# Mapping California’s Urban Forest at Scale: An Error-Adjusted Canopy Time Series for Monitoring Change

**DOI:** 10.64898/2026.05.04.722774

**Authors:** Camille C. Pawlak, Jenn Yost, Jonathan Ventura, Griffen Guizan, Sara Arnold, Gregory Okin, Kyle Cavanaugh, G. Andrew Fricker, Matt Ritter, Thomas W. Gillespie

**Affiliations:** Department of Geography, University of California, Los Angeles, CA, USA; Department of Biological Sciences, Cal Poly, San Luis Obispo, CA, USA; Department of Computer Science & Software Engineering, Cal Poly, San Luis Obispo, CA, USA; Department of Social Sciences, Cal Poly, San Luis Obispo, CA, USA

## Abstract

Statewide tracking of urban tree canopy change is essential for evaluating progress toward policy targets, but detecting real change requires both high-resolution mapping and rigorous uncertainty estimation. We produced a four-year canopy cover time series for all California census-designated places using 60-cm NAIP aerial imagery and a U-Net deep learning model trained with semi-automated LiDAR-derived labels and manually annotated tiles. Canopy cover and change were estimated using stratified, error-adjusted area estimation, enabling comparisons across years. Statewide canopy cover showed a modest negative trend from 2016 to 2022 (Sen’s slope: –0.60% per year), but confidence intervals included zero across all groups and climate zones, indicating that trends were not statistically distinguishable from no change. Urban canopy cover was consistently lower than non-urban canopy by approximately six percentage points, and canopy cover was highest in the Northern California Coast and lowest in the Southwest Desert. Residential parcels accounted for 55-56% of canopy within incorporated urban areas across all years, indicating that statewide canopy increase goals will require engagement with private landowners. Error adjustment substantially altered canopy estimates relative to raw pixel-count totals, with direct implications for AB 2251 canopy tracking where baselines and targets drawn from unadjusted maps may not reflect true canopy extent. This open-source workflow is transferable to future NAIP acquisition years and other U.S. states, providing a scalable framework for long-term urban forest monitoring.

## 1. Introduction

California has committed to increasing urban tree canopy, but measuring progress toward that goal is not straightforward. Even high-resolution canopy maps are predictions, and small apparent changes in canopy cover can fall within the range of classification error, acquisition variability, or image misalignment (Olofsson et al., 2014; Richardson & Moskal, 2014; Lehrbass & Wang, 2012). This creates a central challenge for urban forest monitoring: canopy targets are often expressed as simple percent increases (Morgenroth et al., 2025), while the datasets used to evaluate them rarely propagate uncertainty into change estimates.

Urban tree canopy provides important ecological and social benefits, including cooling, carbon sequestration, reductions in air pollution and stormwater runoff, and associations with physical and mental health (Cavender-Bares et al., 2022; Giacinto et al., 2021; Hwang et al., 2023; Livesley et al., 2016). Over the past decade, California’s urban tree canopy has been affected by storms, extreme droughts, heat events, and expanding urban development (California Department of Conservation, 2020; Hulley et al., 2020; Summers et al., 2022; Williams et al., 2022). Within this context, California’s recent AB 2251 requires a statewide 10% increase in canopy cover by 2035 (California Urban Forestry Act, 2022). Tracking progress toward that goal requires monitoring approaches that can distinguish real canopy gains and losses from apparent change caused by mapping error and image acquisition differences.

High-accuracy canopy maps are increasingly created using high-resolution optical imagery, LiDAR data, and deep learning, often with reported accuracies in the 90-97% range (Akın et al., 2023; Guo et al., 2023; Martins et al., 2021; Timilsina et al., 2020; Velasquez-Camacho et al., 2023; Wang et al., 2021; Zhang et al., 2022). Aerial imagery from programs such as the National Agricultural Imagery Program (NAIP) provides sub-meter imagery statewide and can support fine-scale canopy monitoring. However, canopy cover maps are still often produced at the city scale or for single time points, and existing multi-year analyses often rely on mapped pixel totals without propagating classification error into area estimates. This is especially important when canopy goals are small. In some cities, canopy increase targets are as small as 4% (Morgenroth et al., 2025), meaning apparent change may be similar in magnitude to mapping uncertainty.

To track urban canopy change in California, we produced a four-year (2016, 2018, 2020, 2022) 60-cm canopy cover time series for all California census-designated places (CDPs) and estimated canopy cover and change using error-adjusted areas. This research develops a scalable, open-source workflow for creating canopy cover maps and uses that methodology to answer three primary research questions: (1) How did canopy cover across California CDPs change from 2016 to 2022? (2) How do canopy levels and trends vary across climate zones and between urban and non-urban portions of CDPs? and (3) Within incorporated urban areas, how is canopy distributed across parcel/property types, and is that distribution stable through time? All code, training data, model checkpoints, and canopy raster products are publicly available (GitHub: https://github.com/camipawlak/canopy_cover; Dryad: 10.5061/dryad.rjdfn2zs9).

## 2. Methods

### 2.1 Study Site

California spans an area of approximately 423,970 km^2^ and is home to over 39 million people, with 94% of the population residing in urban areas (U.S. Census Bureau, 2020b). The state is divided into six distinct climate zones: Southwest Desert, Northern California Coast, Southern California Coast, Inland Valleys, Inland Empire and Interior West (McPherson et al., 2010, Figure 2). These zones vary in temperature, precipitation, and vegetation. This study focuses on all CDPs in California, which collectively cover an area of 39,032 km^2^ (U.S. Census Bureau, 2020a; Figure 2). CDPs are populated areas identified by the U.S. Census Bureau for statistical purposes (U.S. Census Bureau, 2016; U.S. Census Bureau, 2018; U.S. Census Bureau, 2020a; U.S. Census Bureau, 2022). For each year, the extent of the raster data is the extent of that year’s CDP (2016: 37,911.90 km², 2018: 38,014.60 km², 2020: 39,032.35 km², 2022: 39,074.13 km²). Unlike many previous datasets and studies that concentrate only on urban areas (CAL FIRE, 2025; Kropp, 2024; MacFaden et al., 2012; Zhang et al., 2022), this research also includes less urbanized and non-urban CDPs.

### 2.2 Approach

We conducted a two-step process to generate an urban canopy cover map for all census-designated places in California. First, we developed a general model by creating automated training data using LiDAR-derived canopy height models (CHMs) and NAIP imagery. A U-Net (Ronneberger et al., 2015) neural network was trained on these data, with training data stratified to match the proportion of urban areas in each climate zone. Second, we used transfer learning (Bozinovski and Fulgosi et al., 1976) to fine-tune the LiDAR based model once for each climate zone and statewide using manual annotated training data. The best fine-tuned model was applied to the CDPs within each climate zone across California using imagery from 2016, 2018, 2020, and 2022. Mapped canopy areas were error-adjusted using accuracy assessments (Figure 1).

**Figure 1.**
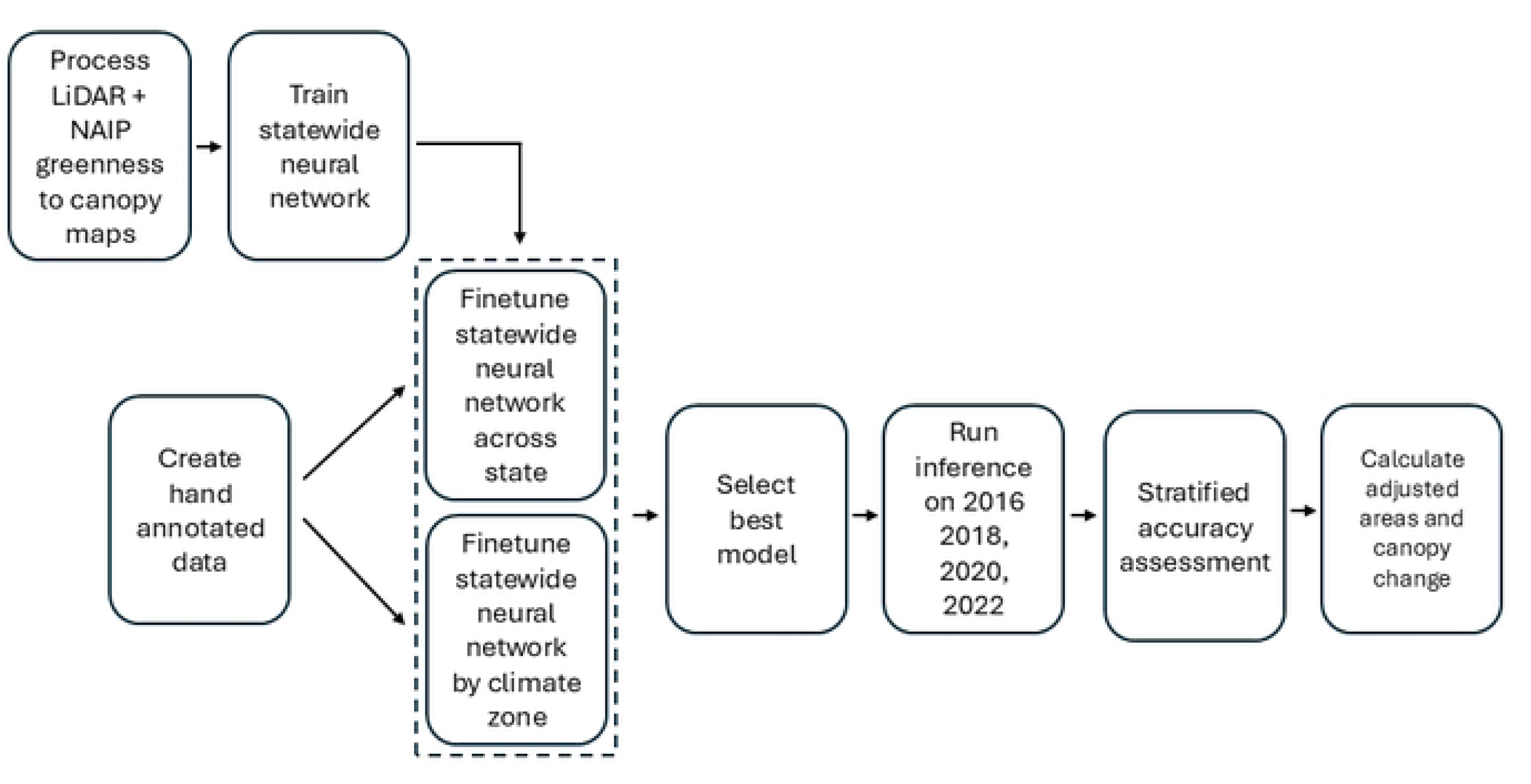
Workflow for generating California canopy cover maps and estimating canopy change from 2016 to 2022. LiDAR-derived canopy height models were paired with NAIP imagery and an NDVI-based filter to create semi-automated canopy masks used to train a statewide U-Net model. Manually annotated tiles were used for two fine-tuning strategies: (1) statewide fine-tuning across all climate zones and (2) climate-zone-specific fine-tuning. The best-performing model was selected and applied to NAIP imagery from 2016, 2018, 2020, and 2022. A stratified reference sample was used to assess accuracy and to compute error-adjusted canopy area estimates and confidence intervals for canopy cover and change.

### 2.3 Data Sources

U-Net inputs were NAIP multispectral imagery, collected approximately every two years. In California, NAIP imagery has been available at 60 cm resolution since 2016. LiDAR data was sourced from the USGS 3D Elevation Program (3DEP) and included datasets for 28 cities across California from 2016 to 2021 (Figure 2, Appendix C). LiDAR datasets varied in density and coverage, ranging from 3.47 – 38.44 points per m^2^.

**Figure 2.**
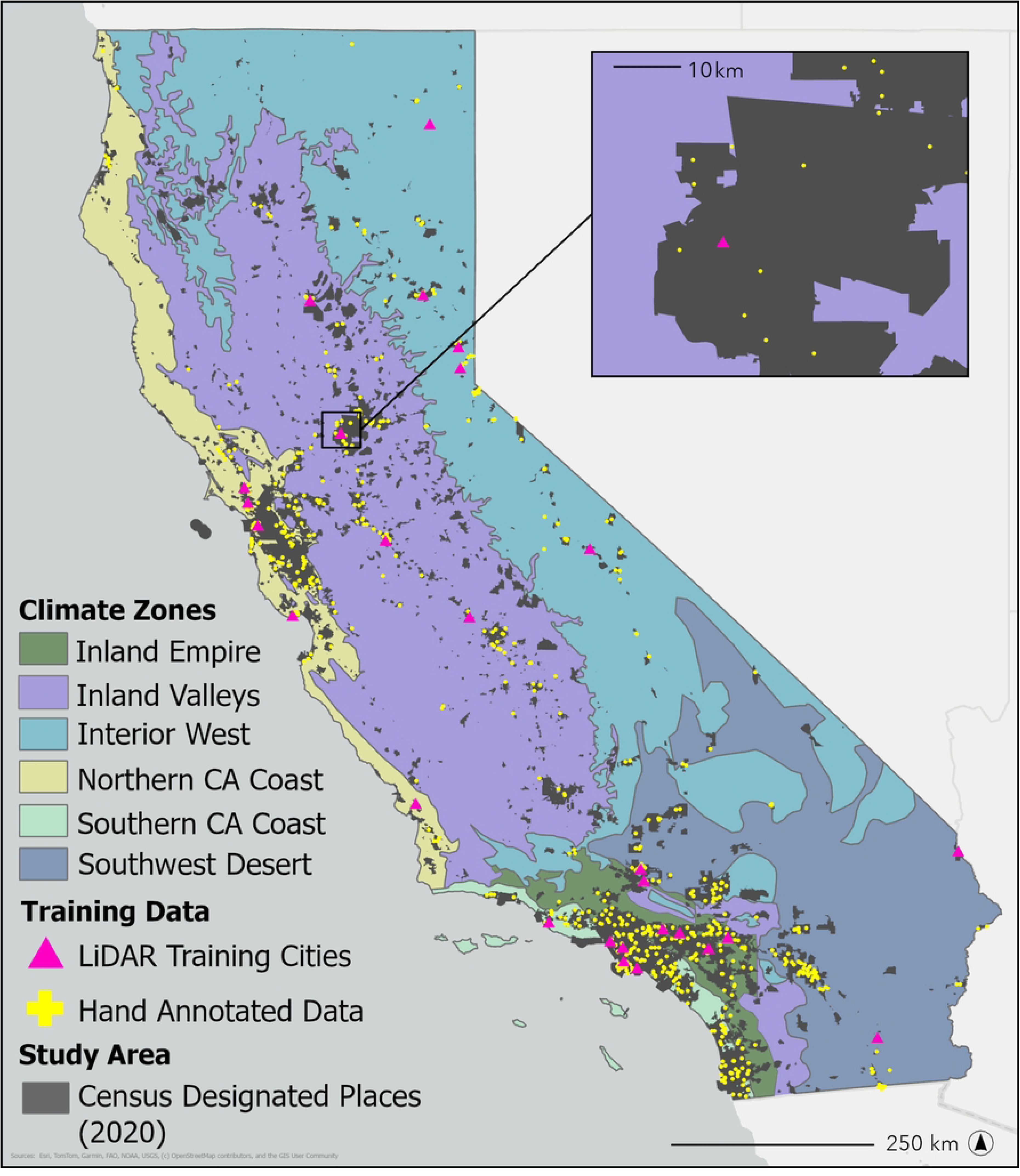
The climate zones of California, CDPs (2020), and locations of LiDAR training cities and manually annotated tiles.

### 2.4 LiDAR Generated Training Data

In the first stage of model development, we created a semi-automated canopy cover label dataset using USGS LiDAR data. Previous studies (Weinstein et al., 2019) have demonstrated the efficiency of automated training data for canopy cover prediction, which informed our methods.

Cities were selected to provide training data to represent typical urban areas within each of California’s six climate zones (Figure 1, Appendix C). For each selected city, we processed the LiDAR point cloud data using the Point Data Abstraction Library (PDAL) to generate Digital Surface Models (DSM), Digital Terrain Models (DTM), and Canopy Height Models (CHM). DSMs were created by filtering first-return points to capture surface elevations, including vegetation and structures, and interpolating the data using Inverse Distance Weighting. DTMs were generated by isolating ground points and interpolating bare-earth elevations using Delaunay triangulation to create a continuous surface. The CHM was calculated by subtracting the DTM from the DSM.

The resulting CHMs were paired with NAIP imagery from the nearest available year (all data within one year of the NAIP flyover). We classified all areas with heights ≥ 1.8 m as canopy (value = 1) and other areas as non-canopy (value = 0). Because urban areas include structures such as buildings, which were often misclassified as canopy, we refined the labels using Normalized Difference Vegetation Index (NDVI) values derived from NAIP imagery. Pixels with NDVI ≤ 0.05 were reclassified as non-canopy. This threshold was selected based on visual inspection of training data across multiple cities, where values at or below this level consistently corresponded to non-vegetated surfaces such as pavement, rooftops, and bare soil, with no clear cases of tree canopy falling below this value. This semi-automated process produced 30,458 training masks (448 ×448 pixels) across the state, reducing the time and cost compared to manual annotation.

### 2.5 Manually Annotated Training Data

We created 645 manually annotated tiles from 2020 NAIP imagery to fine-tune models statewide and by climate zone (Figure 2). Tiles were 153.6 × 153.6 m, or 256 × 256 pixels. Initial tile locations were selected using stratified random sampling by climate zone, with allocation proportional to CDP area. Additional tiles were added iteratively to improve regional representation, resulting in 83-134 tiles per climate zone. Annotators delineated tree canopy using true-color and false-color infrared imagery. Uncertain areas, especially tree-shrub boundaries, were checked using Google Street View and visual cues such as shadows, crown structure, and canopy overlap with buildings. Each tile was annotated by one person and reviewed by a second annotator for consistency. The goal of annotations was to only include canopy over 1.8 m in height as best as possible using visual imagery.

### 2.6 Imagery Pre-Processing

The NAIP imagery inputs consisted of four spectral bands: red, green, blue, and near-infrared, with a resolution of 60 cm as well as a fifth band, the Normalized Difference Vegetation Index (NDVI), to help make vegetation stand out. NDVI was calculated using the formula:

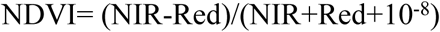

The small constant 10^-8^ was included to prevent division by zero. NDVI values, which theoretically range from −1 to 1, were linearly scaled to 0-255 by applying the transformation (NDVI + 1) × 127.5, mapping −1 to 0 and +1 to 255. The four NAIP spectral bands were then independently normalized to 0-255 using minimum-maximum normalization, with bounds pre-computed from the full California NAIP dataset for 2020 and applied as fixed values (red: 0-222, green: 0-221, blue: 0-224, NIR: 0-216). The processed data, including all five bands, was divided into 448 ×448 pixel tiles for those paired with LiDAR generated masks, and 256 ×256 pixel tiles for the manually annotated data. These tiles were split into training (80%), testing (10%), and evaluation (10%) datasets for model development.

### 2.7 Model Architecture and Training

A convolutional neural network with a U-Net architecture (Ronneberger et al., 2015) was chosen due to its ability to handle complex decision boundaries, particularly in distinguishing between trees, shrubs, and grass patches, which can have similar visual characteristics in aerial imagery. U-Net has been widely used in segmentation tasks, including tree canopy segmentation (Martins et al., 2021; Ronneberger et al., 2015; Wang et al., 2021), and has consistently outperformed or matched other deep learning architectures for tree canopy segmentation.

The model used a U-Net encoder-decoder architecture with skip connections and a ResNet-34 backbone (He et al., 2015). The decoder used four transposed-convolution blocks, and the final 1 × 1 convolution produced a per-pixel canopy probability map. These probabilities were converted to a binary output by applying a threshold of 0.65, selected based on visual inspection of validation outputs where this value balanced commission and omission errors in canopy classification. Values above the threshold were classified as canopy (1) and those below as non-canopy (0). To handle no data regions, areas where all input bands were zero were excluded and assigned a no data value (255).

A masked binary cross-entropy loss function was used to optimize the model, applied to logits directly with pixels assigned a no-data value of 255 excluded from the loss calculation. The model was implemented using PyTorch and trained with the AdamW optimizer (Loshchilov and Hutter, 2019), with a learning rate of 0.0001 and a batch size of 16. Training was conducted on two Tesla V100 GPUs.

The SLM was trained for 150 epochs on 24,366 LiDAR-derived training chips, with the best checkpoint saved based on validation loss. This checkpoint was then used as the starting point for two fine-tuning runs. The SFT was trained for 147 epochs on 516 manually annotated tiles drawn from across all climate zones. The RFT produced one model per climate zone, each trained on region-specific manually annotated tiles; training set sizes ranged from 65 to 107 tiles per zone and the number of epochs at the best checkpoint varied by zone (14 – 89 epochs), reflecting differences in dataset size and convergence rate (Table 1). All fine-tuning used end-to-end training with data augmentation including random 90° rotations (0°, 90°, 180°, 270°) and random horizontal flips. For all models, the best checkpoint was selected based on validation loss, alongside visual inspection of inference outputs to catch spatial artifacts or boundary errors not captured by loss alone.

**Table 1:**
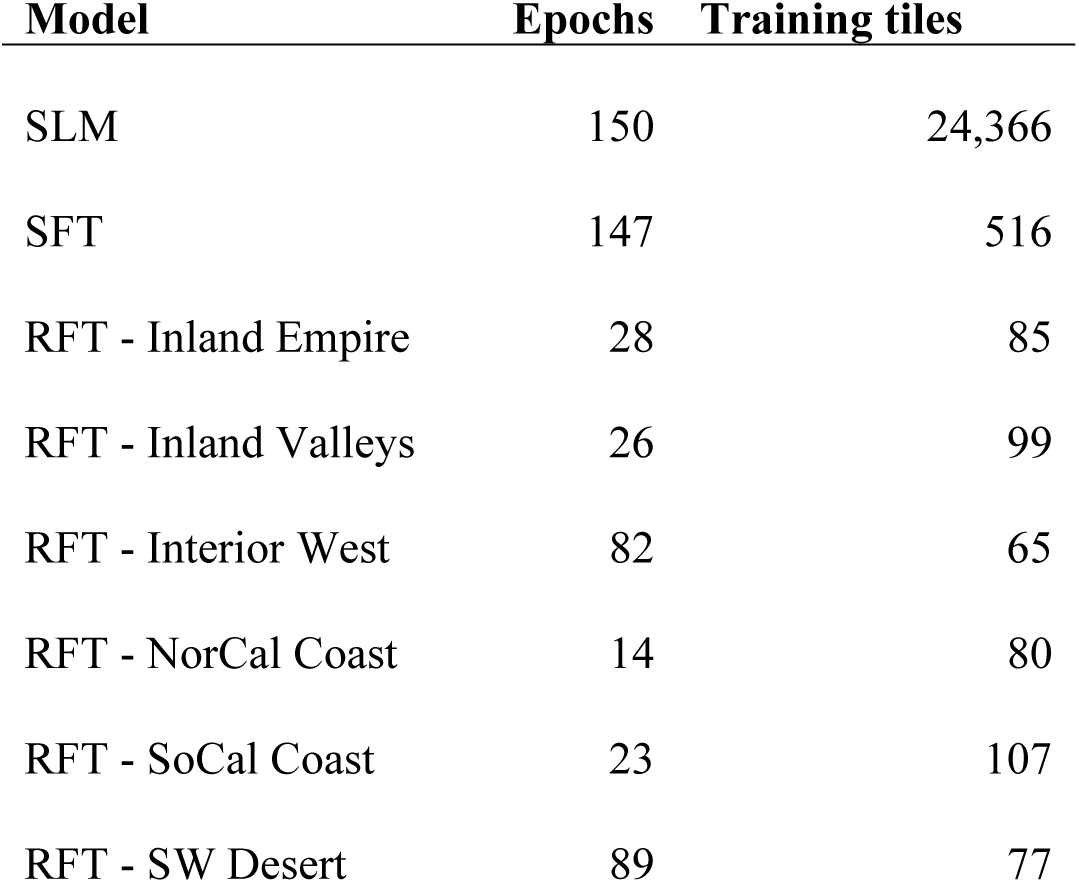
Number of epochs used to train selected versions of models, and number of tiles per model used for training (after 80/10/10 split for training, testing, and validation).

### 2.8 Accuracy Analysis and Area Estimation

We evaluated 2020 model performance using user’s accuracy, producer’s accuracy, and F1 scores on manually annotated data withheld from training. Validation was assessed on both the statewide and fine-tuned models. To assess the accuracy of change over time and accuracy of each year’s output, we took a stratified random sampling approach based on Olofsson et al. (2014) with the classes “canopy” and “non-canopy” as classification strata, and “urban” and “non-urban” (in 2020) as geographic strata. Urban areas were defined using yearly NLCD products developed areas (Dewitz et al., 2019). Within each CDP, we converted areas of low, medium, and high developed intensity to a polygon, then filled all holes smaller than 1 km^2^ to ensure urban parks were included as “urban.” The remainder of the CDPs were classified as non-urban. Prior to adjustment of area, we summed mapped canopy area into groups by assigning CDPs into urban and non-urban areas, then assigning climate zones to each urban and non-urban section of the CDP. We removed all water bodies and coastline from the CDPs prior to total area calculations (areas listed in section 2.1 are for CDPs are prior to water removal). The reference sample was placed using 2020 urban boundaries; for area estimation in each year, urban and non-urban areas were re-defined using that year’s NLCD product to compute year-specific mapped-area weights.

Initially, 1,200 points were randomly placed within 2020 urban areas (600 on mapped canopy, 600 on mapped non-canopy). Then 500 points were placed in non-urban areas with a 50/50 split for mapped canopy and non-canopy. Each sample point was assigned a reference classification based on the pixel in which it fell; pixels were labeled by a human as canopy or non-canopy based on the dominant cover of the full pixel extent. Ambiguous pixels were labeled as NA and excluded from the analysis. Each point was validated once per year against NAIP imagery (4,800 unique validations were completed in this process). Unclear points were checked with Google Street View where available and classified as NA if uncertainty remained. This most frequently happens at the edge of a tree canopy, on shrubs where height could not be estimated, or in very dark shadows. To ensure accuracy could be assessed by climate zone, additional points were generated using simple random sampling (Python random module) for canopy and non-canopy within each climate zone to ensure no class was too rare, following Olofsson et al. (2014), who recommend increasing sample sizes for rare classes to support meaningful accuracy estimates. A minimum of 50 points per canopy and non-canopy class per climate zone was created (Appendix A).

To estimate canopy area for each year, we applied stratified, error-adjusted area estimation following Olofsson et al. (2014) to our mapped values. For each year, we defined two map strata based on the classified raster: canopy (mapped class 1) and non-canopy (mapped class 0). Within each reporting domain (statewide, urban vs non-urban, and climate zone), we calculated the mapped-area weight for each stratum as:

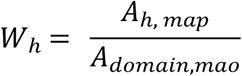

where A_h,map_ is the mapped area in stratum *h* and A_domain,map_ is the total mapped area in that domain.

We then summarized the reference sample into error matrices for canopy and non-canopy in each domain. Reference NAs were counted for reporting but excluded from all further estimation steps, only reference points of 0 and 1 were used for accuracy calculations. Error-matrix cell counts were converted to estimated area proportion using a stratified estimator:

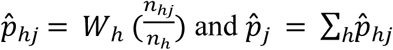

where *n_hjj_* is the count of reference samples in class *j* within the mapped stratum *h*, and *n_h_* is the total number of reference points in that stratum. Overall accuracy, user’s accuracy, and producer’s accuracy were computed from the estimated error matrix proportions, standard errors and 95% confidence intervals for area proportions and error-adjusted areas were calculated using the stratified variance estimators in Olofsson et al. (2014) (Appendix A). This approach corrects for classification error that would otherwise cause pixel-count totals to over– or underestimate true canopy area. Using adjusted areas, we compared statewide canopy percent across years, across urban and non-urban area, across climate zones, and across urban and non-urban areas within climate zones.

To understand potential drivers of canopy change, we also summed canopy area by parcel type to assess where canopy change is happening over time. Parcel data and classification of property type are only for incorporated urban areas and the layer is from 2013 (McPherson et al., 2017). For each parcel type (Residential (Muti-family, MF), Residential (Single Family, SF), Commercial/Industrial/Institutional, Open Space, Transportation Corridors, Water, and Unclassified) we built an error matrix from points in that parcel class, again excluding NAs, estimated the probability that mapped canopy and non-canopy were truly canopy, and applied the Olofsson et al. (2014) methods of adjustment described above to estimate true canopy area. We then converted our areas to percent of statewide canopy in each parcel type, with uncertainty summarized as 95% intervals (Appendix B).

### 2.9 Change Analysis

We compared adjusted canopy area estimates across the time series statewide, in urban and non-urban areas, in each climate zone, and within urban and non-urban areas in each climate zone, and between adjacent years in each climate zone. Differences (for example, urban minus non-urban within a year) were evaluated with a Wald Z-test using the estimated difference in error-adjusted canopy cover divided by the standard error of that difference. We used two-sided tests with α=0.05. To assess change over the broader time series, we calculated Sen’s slope using the median of all pairwise slopes between years. Uncertainty bounds for Sen’s slope were approximated using the empirical distribution of pairwise slopes (2.5^th^-97.5^th^ percentiles). Sen’s slope was computed from error-adjusted canopy estimates.

## 3. Results

### 3.1 Model Accuracy 2020

Using 2020 NAIP manually annotated data withheld from the training dataset, we compared model accuracy of the statewide LiDAR model (SLM), the regionally fine-tuned model (RFT), and the statewide fine-tuned models (SFT) across climate zones. In all cases, the SFT model had the highest F1 score and producer’s accuracy (Table 2). User’s accuracy was also the highest for SFT, except in the case of Inland Valleys, where it was slightly lower than the RFT model (Table 2).

**Table 2:** User’s accuracy, producer’s accuracy, and F1-score comparison for the Statewide LiDAR Model (SLM), Regional Fine-Tuned Model (RFT), and Statewide Fine-Tuned Model (SFT). The metrics demonstrate the performance of each model in predicting canopy cover, with RFT capturing regional specificity, SFT benefiting from broader manually annotated data, and SLM serving as the baseline trained on statewide LiDAR-derived data. These numbers are based on 2020 imagery.

| **Region** | **Model** | **User's** | **Producer's** | **F1** |
| --- | --- | --- | --- | --- |
| *Southern California Coast* | SLM | 0.80 | 0.68 | 0.73 |
|  | RFT | 0.89 | 0.76 | 0.82 |
|  | SFT | **0.91** | **0.83** | **0.87** |
| *Inland Empire* | SLM | 0.82 | 0.75 | 0.78 |
|  | RFT | 0.90 | 0.77 | 0.83 |
|  | SFT | **0.91** | **0.86** | **0.88** |
| *Northern California Coast* | SLM | 0.89 | 0.60 | 0.72 |
|  | RFT | 0.91 | 0.77 | 0.84 |
|  | SFT | **0.93** | **0.87** | **0.90** |
| *Inland Valleys* | SLM | 0.89 | 0.53 | 0.67 |
|  | RFT | **0.92** | 0.81 | 0.86 |
|  | SFT | 0.91 | **0.87** | **0.89** |
| *Interior West* | SLM | 0.80 | 0.71 | 0.75 |
|  | RFT | 0.86 | 0.84 | 0.85 |
|  | SFT | **0.92** | **0.88** | **0.90** |
| *Southwest Desert* | SLM | 0.77 | 0.71 | 0.74 |
|  | RFT | 0.89 | 0.75 | 0.82 |
|  | SFT | **0.95** | **0.87** | **0.91** |

The highest statewide fine-tuned F1 score was for the Southwest Desert (F1 = 0.91) and the lowest for the Southern California Coast (F1=0.87) (Table 2). Among regional models, the highest F1 was for the Inland Valleys (F1=0.86), and the lowest F1 was a tie for the Southern California Coast and Southwest Desert (F1=0.82) (Table 2). Figure 3 shows example model outputs.

**Figure 3.**
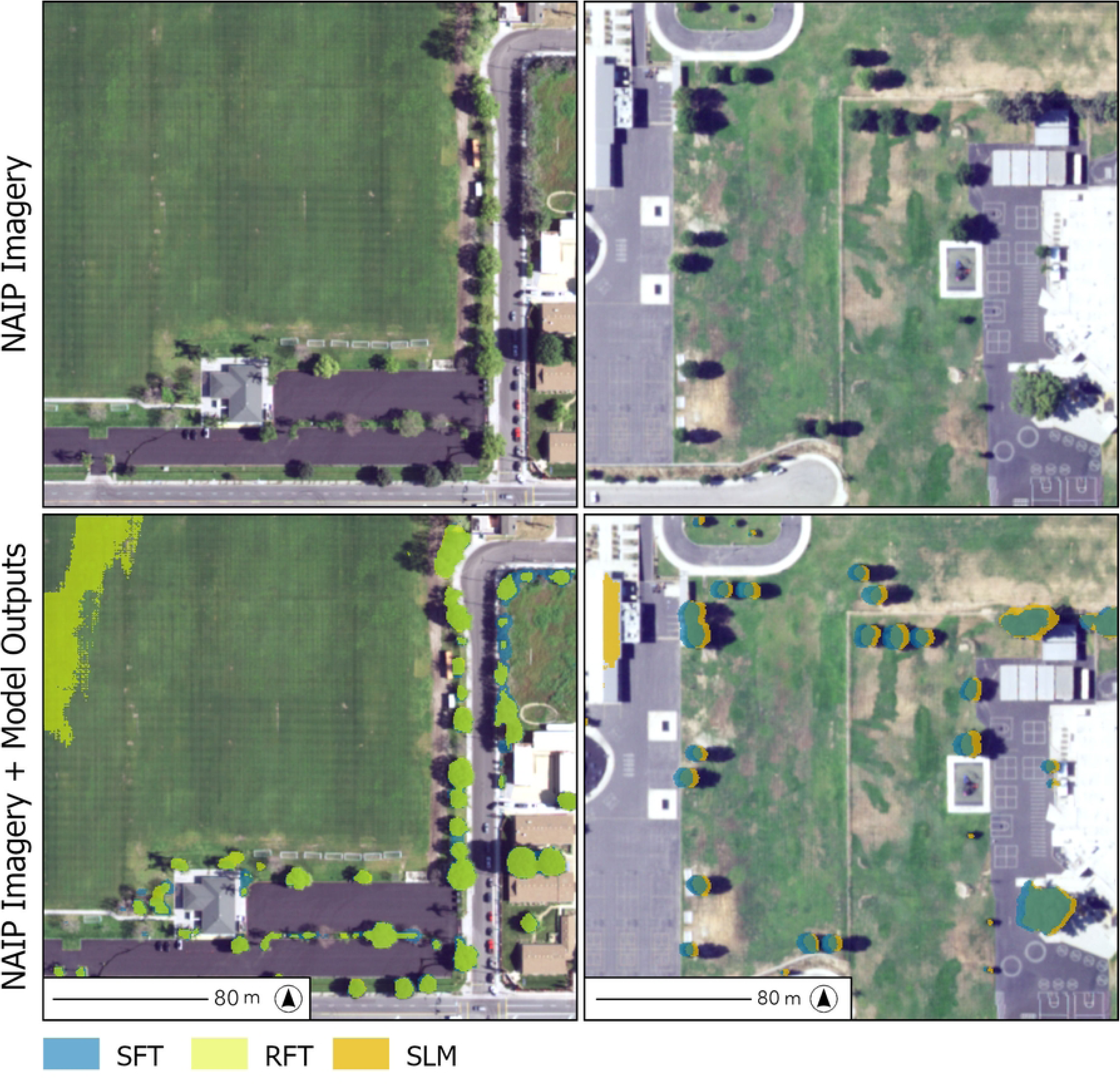
Example model outputs for canopy segmentation on NAIP imagery. The top panel shows the NAIP image tile with a corresponding bottom panel with transparent model outputs overlaid together. Left: The statewide fine-tuned model (blue) overlaid under the regionally fine-tuned model (yellow). RFT has more artifacts, and under predicts canopy, but is sometimes more precise than SFT. Right: SFT (blue) overlaid on top of the output from the statewide LiDAR model (SLM, orange). SLM produces artifacts on rooftops, and tends to overpredict canopy in shadows and beyond the extent of the tree compared to SFT.

### 3.2 Extrapolation to other years

Model accuracy was the highest for 2020, the year training data was created for. However, model accuracy remained high across the three other applied years with overall accuracy at 0.90 and above for all years (Table 3).

**Table 3.**
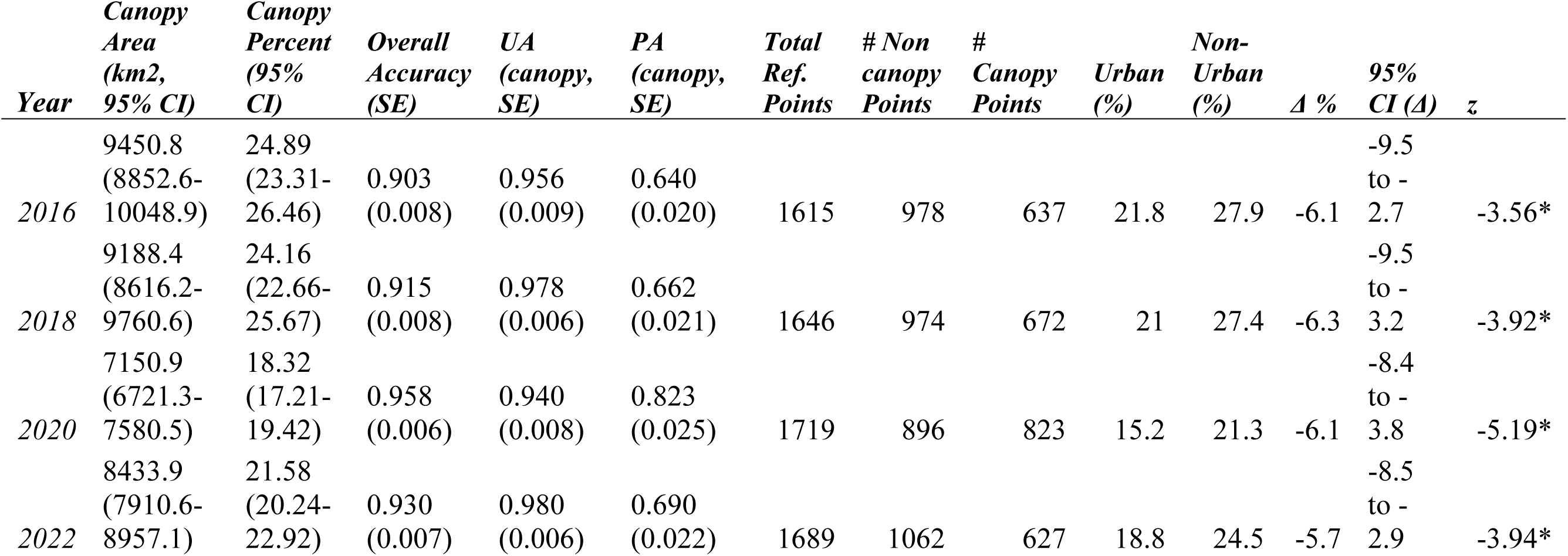
Error-adjusted canopy area and cover estimates for each year across all California CDPs, with accuracy metrics and urban/non-urban comparisons. Canopy area (km²) and canopy cover percent are reported with 95% confidence intervals. Overall accuracy, user’s accuracy (UA), and producer’s accuracy (PA) for the canopy class are reported with standard errors and are based on total reference points (excluding NAs) for that year. Urban and non-urban canopy cover percentages are error-adjusted estimates. Δ reports the difference in canopy cover percentage points between urban and non-urban areas within each year, with 95% confidence interval, Wald Z statistic, and p-value from a two-sided test marked with * for below 0.01.

### 3.3 Change across California

Error adjusted statewide canopy cover was 24.9% in 2016 and 21.6% in 2022. Across all years, error-adjusted canopy cover was lower in urban than non-urban areas. Urban versus non-urban canopy cover was 21.8% vs 27.9% in 2016, 21.0% vs 27.4% in 2018, 15.2% vs 21.3% in 2020 and 18.8% vs 24.5% in 2022 (Table 3, Figure 4). Across the full time series, Sen’s slope indicated a modest decline in canopy cover statewide (–0.60% per year), with similar rates of change in urban (–0.53% per year) and non-urban areas (–0.64% per year). However, confidence intervals for all three estimates were wide and included zero, indicating that these trends are not statistically distinguishable from no change over this period (Table 4).

**Figure 4.**
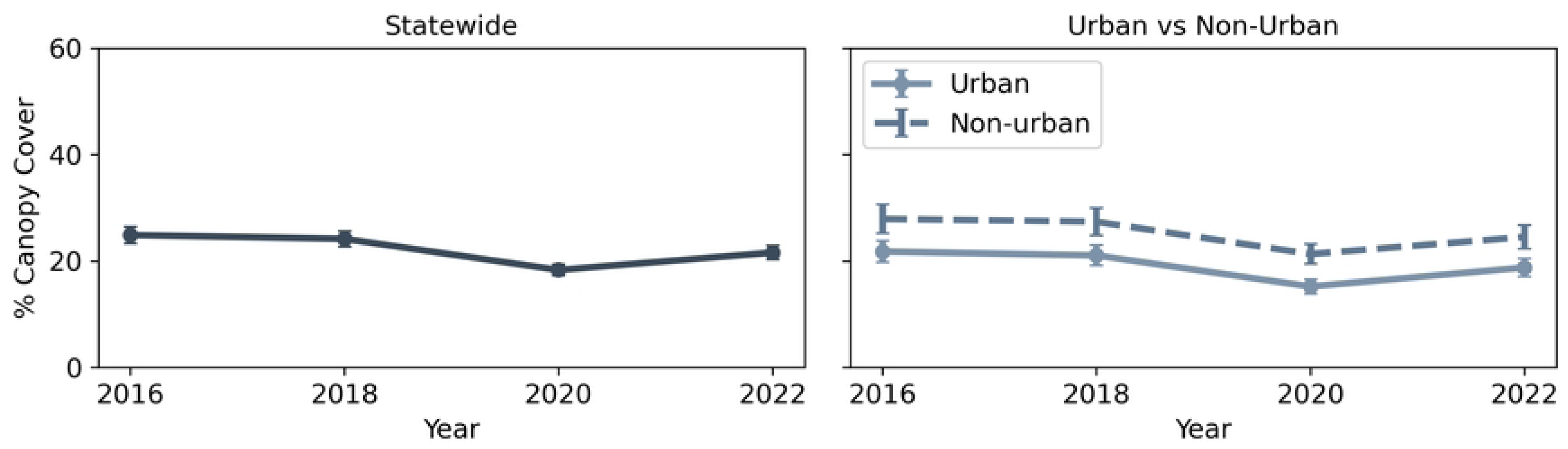
Error-adjusted canopy cover (%) across California census-designated places (CDPs) from 2016 to 2022. The statewide time series (left) summarizes canopy cover across all CDPs, and the time series (right) separates CDP area into urban and non-urban classes. Points show error adjusted canopy percent. Error bars indicate 95% confidence intervals (estimate ± 1.96 x SE).

**Table 4:** Sen’s slopes calculated using error-adjusted area estimates across the time series statewide, for urban and non-urban areas, and in each climate zone. All confidence intervals cross zero, suggesting that trends are not statistically distinguishable from no change over the study period.

| *Group* | *Sen’s*<br>*Slope* | *CI low* | *CI high* |
| --- | --- | --- | --- |
| *Statewide* | -0.60 | -2.76 | 1.38 |
| *Urban* | -0.53 | -2.77 | 1.52 |
| *Non-Urban* | -0.64 | -2.86 | 1.36 |
| *Inland Empire* | -0.55 | -3.05 | 3.46 |
| *Inland Valleys* | -1.15 | -3.10 | 0.51 |
| *Interior West* | -0.55 | -2.56 | 1.49 |
| *Northern California Coast* | -0.53 | -1.05 | 0.48 |
| *Southern California Coast* | -0.90 | -4.06 | 1.96 |
| *Southwest Desert* | 0.18 | -2.01 | 2.20 |

When evaluated across sequential time intervals, significant changes in canopy cover were concentrated in the 2018-2020 period. During this interval, canopy cover declined significantly in the Inland Empire (–6.38%), Inland Valleys (–6.58%), and Southern California Coast (–8.60%) (Wald Z test, p < 0.01). No climate zones exhibited significant changes between 2016 and 2018, and only limited recovery was observed between 2020 and 2022, with a significant increase detected in the Inland Empire (+7.80%). All other changes were not statistically distinguishable from zero (p > 0.05).

Across climate zones, the Northern California Coast had the highest average canopy cover percent across four years at 29.7%, then Inland Valleys (27.9%), Interior West (27.6%), Southern California Coast (18.2%), Inland Empire (16.3%), and finally Southwest Desert at 7.1% (Table 5, Figure 5).

**Figure 5.**
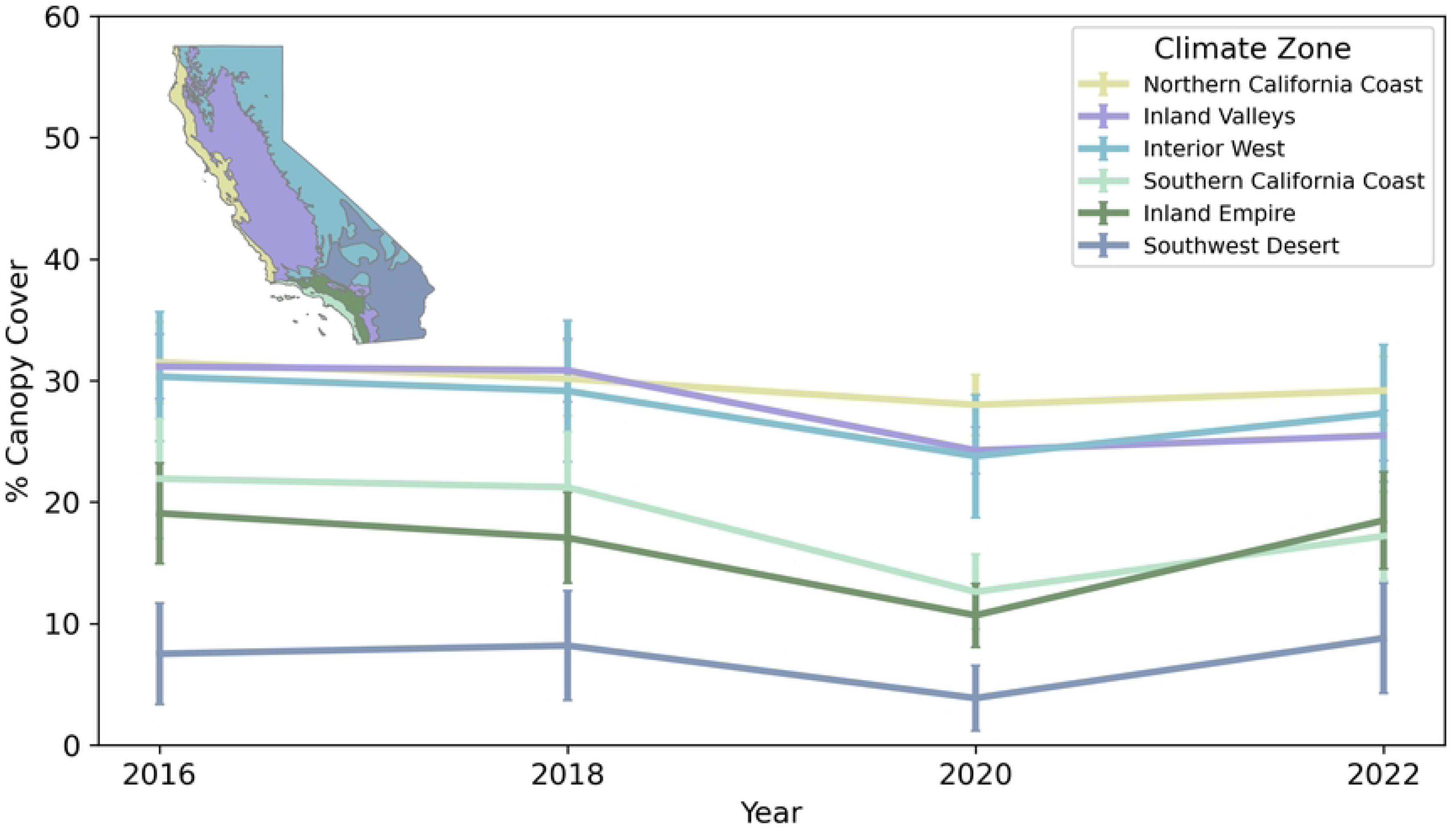
Error-adjusted canopy cover (%) by California climate zone from 2016 to 2022. Error bars indicate 95% confidence intervals (estimate ± 1.96 x SE). Climate zones are shown on the upper left map.

**Table 5.** The percent of CDP area in each climate zone, as well as the share (reported as percent) of total state canopy within CDPs that occurs in each climate, and the percent canopy cover within each climate zone in California. Values are reported based on 2020 only (the highest accuracy classification).

| *Climate Zone* | *% of Total*<br>*State CDP*<br>*Area* | *% of Total*<br>*State CDP*<br>*Canopy* | *% Canopy Cover (95% CI)* |
| --- | --- | --- | --- |
| *Inland Empire* | 17.11 | 9.96 | 10.68 (8.07-13.29); SE 1.33 |
| *Inland Valleys* | 33.47 | 44.22 | 24.26 (22.34-26.19); SE 0.98 |
| *Interior West* | 9.8 | 12.69 | 23.78 (18.72-28.84); SE 2.58 |
| *Northern California Coast* | 13.84 | 21.11 | 28.01 (25.52-30.50); SE 1.27 |
| *Southern California Coast* | 13.85 | 9.51 | 12.61 (9.53-15.70); SE 1.57 |
| *Southwest Desert* | 11.92 | 2.51 | 3.87 (1.16-6.57); SE 1.38 |

When evaluated by climate zone, Sen’s slope estimates indicated generally small and uncertain rates of change in canopy cover from 2016 to 2022 (Table 4). Estimated trends were negative in most climate zones, including the Inland Empire (–0.55% per year), Inland Valleys (–1.15% per year), Interior West (–0.55% per year), Northern California Coast (–0.53% per year), and Southern California Coast (–0.90% per year), while the Southwest Desert showed a slight positive trend (0.18% per year). However, confidence intervals for all climate zones included zero (Table 4), indicating that none of these trends were statistically distinguishable from no change over the study period.

Accuracy assessment points were insufficient to compute error-adjusted estimates for urban and non-urban areas within individual climate zones, so mapped canopy values are reported for these comparisons. While mapped values may over– or underestimate true canopy extent, they are sufficient to characterize relative differences across climate zones. Within each climate zone, non-urban areas in the Northern California Coast, Inland Valleys, and Interior West had higher mapped canopy cover than their urban counterparts. The Inland Empire and Southern California Coast showed similar canopy cover in urban and non-urban areas, while the Southwest Desert had lower canopy cover outside of urban areas (Figure 6).

**Figure 6.**
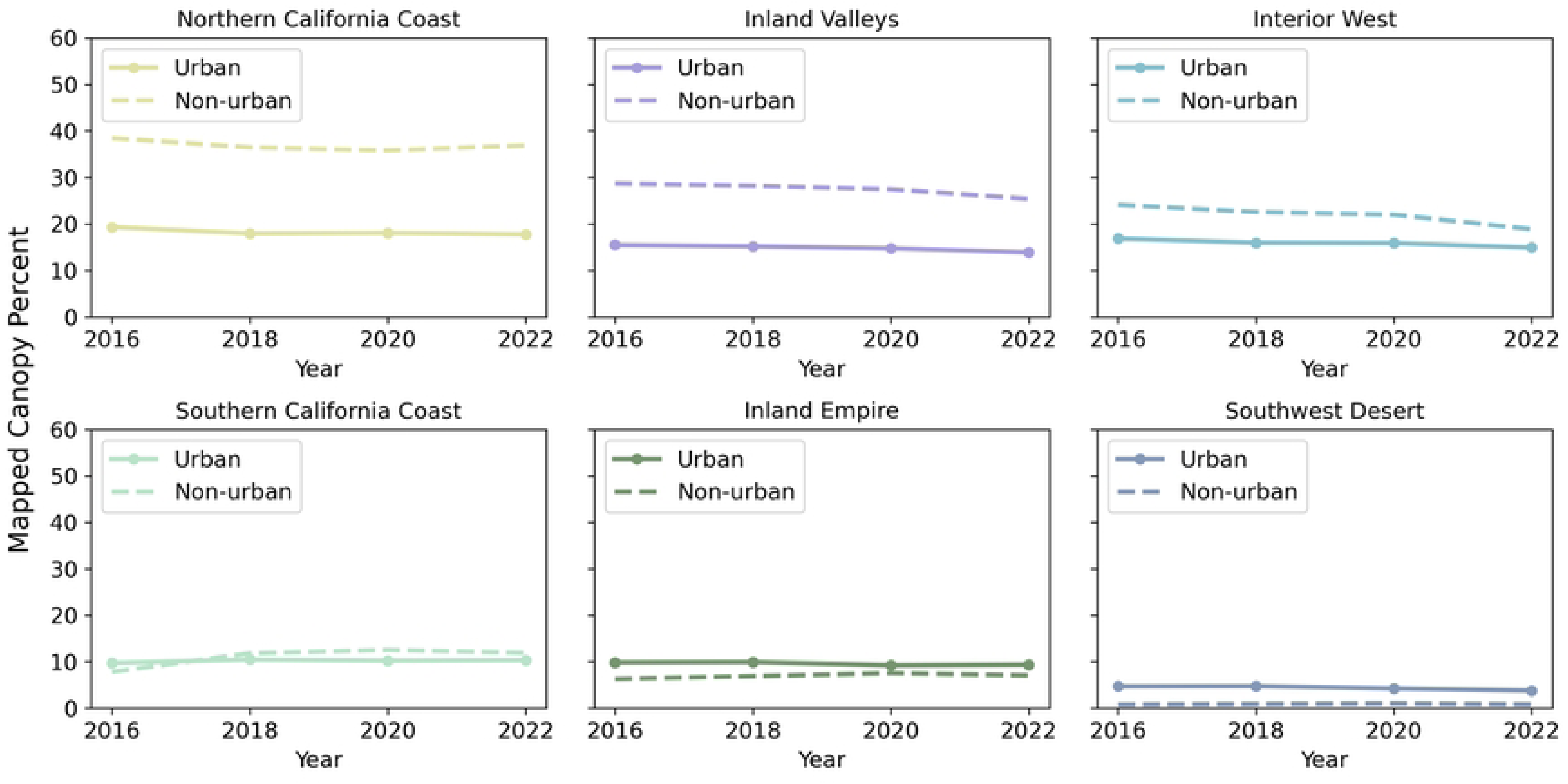
Mapped values for percent canopy cover in each climate zone’s urban and non-urban areas. Most climate zones have higher canopy cover in non-urban areas.

### 3.4 Property Types with Change

The distribution of canopy cover across parcel types was largely stable across years, with each parcel type contributing a similar share of total canopy cover in each year. Residential multifamily and single-family parcels collectively made up between 55% and 56% of the canopy cover each year. Open space accounted for between 17% and 21% each year. Commercial, industrial, and institutional properties made up between 11% and 13% each year. Roadways made up between 11% and 14% each year (Figure 7, confidence intervals in Appendix B).

**Figure 7.**
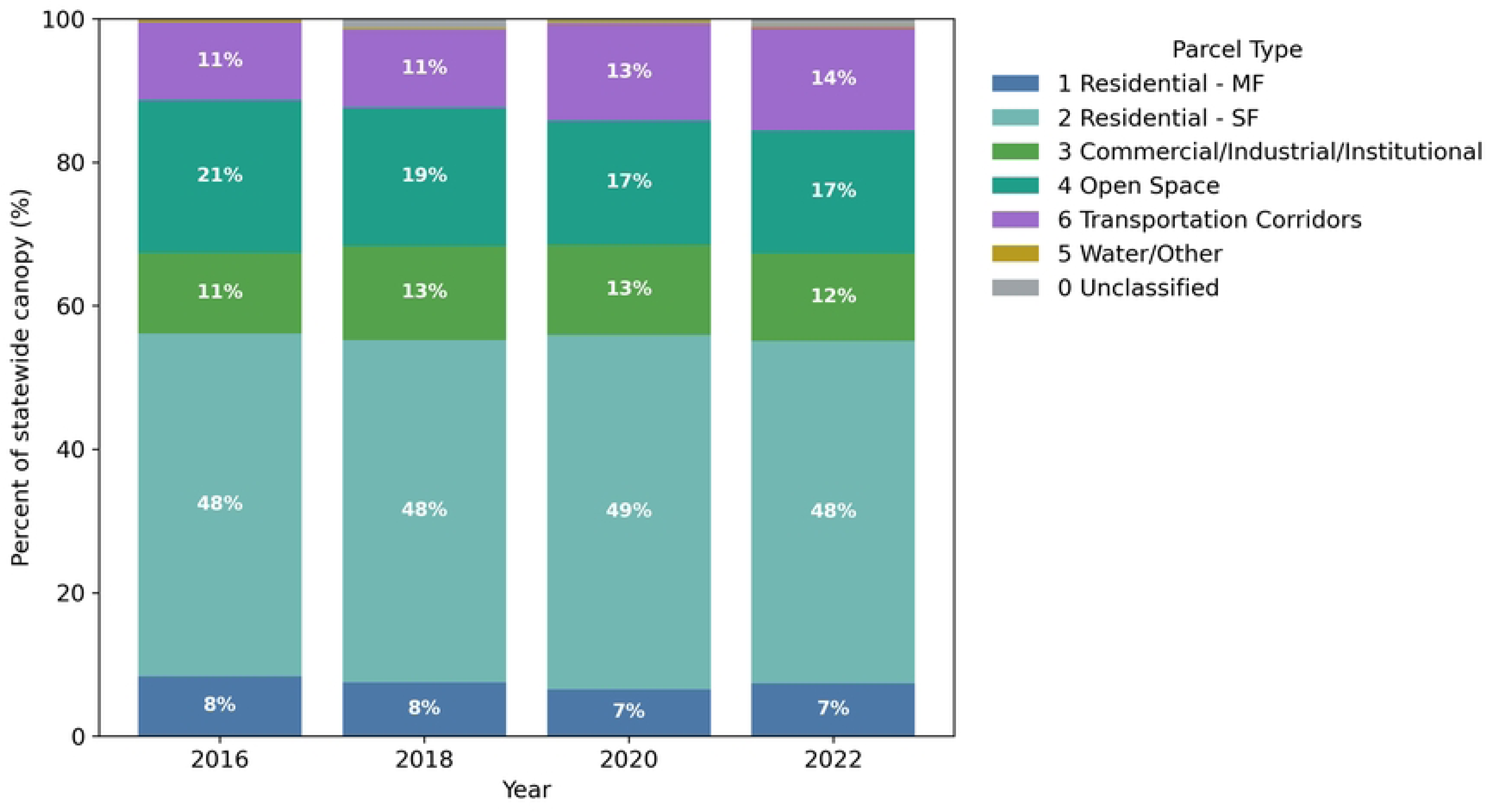
Error-adjusted parcel-type percent share of California statewide canopy by year. Stacked bars show the percent share of statewide adjusted canopy area attributed to each parcel type for 2016, 2018, 2020, and 2022.

## 4. Discussion

This study produces an error-adjusted multi-year urban canopy dataset for California census-designated places across four acquisition years at 60 cm resolution. By combining semi-automated LiDAR-derived labels with targeted manual annotation, we achieved consistently high classification accuracy across years while creating a workflow transferable to future NAIP acquisitions and other states where NAIP and 3DEP LiDAR are available. Canopy area and change were estimated using stratified, error-adjusted area estimation following Olofsson et al. (2014), which we demonstrate produces substantially different estimates than raw pixel counts, a distinction with direct consequences for tracking progress toward AB 2251.

### 4.1 Canopy in California

Statewide canopy cover exhibited an insignificant decline between 2016 and 2022, with the largest decrease occurring between 2018 and 2020 and a partial rebound by 2022. This decline was not uniform across the state; significant decreases during 2018-2020 were concentrated in the Inland Empire, Inland Valleys, and Southern California Coast, and this was the only interval in which statistically significant declines were detected across climate zones. This canopy decrease may reflect a combination of real canopy loss and view angle acquisition variability. Given the two-year NAIP acquisition rate and known variability in aerial imagery geometry, longer time series are required to distinguish persistent structural canopy change from short-term imagery variability. The magnitude of observed change was small relative to total canopy extent, underscoring the importance of uncertainty-adjusted values when interpreting trends.

Spatial patterns broadly reflected California’s climate gradient: canopy cover was highest in the Northern California Coast, Inland Valleys, and Interior West, and lowest in the Southwest Desert. Across all climate zones, canopy cover was consistently lower in urban than non-urban areas, although the magnitude of this varied regionally. In the Southern California Coast, urban and non-urban canopy cover were often similar, whereas in the Inland Empire and Southwest Desert, urban canopy occasionally exceeded non-urban canopy. In these regions, shrub-dominated landscapes are common and are included in our canopy definition (woody vegetation ≥1.8 m), which can reduce contrasts between developed and non-developed areas. Except for mountainous areas and riparian corridors, the Los Angeles Basin was not historically a densely forested ecosystem, so lower canopy cover in this region should not be interpreted as forest loss (Ethington et al., 2020; Pawlak et al., 2023).

Maintaining high canopy cover percentages in urban areas will likely depend less on rapid gains from new plantings and more on preventing losses. Large, mature trees contribute disproportionately to ecosystem services (Doick, 2021; Liang and Huang et al., 2023) and the removal of a single large tree typically cannot be offset quickly by planting multiple small trees (Le Roux et al., 2014), especially within short time periods. Each year, the majority of canopy cover was on privately managed property types. This is especially important because tree canopy is often inequitably distributed: across 5,723 U.S. communities, low-income blocks had 15.2% less tree cover and were 1.5°C hotter than high-income blocks on average (McDonald et al., 2021). Because canopy is concentrated on private residential parcels, and because census tracts with lower income and higher proportions of non-white residents tend to have fewer and slower-growing large street trees (Velasquez-Camacho et al., 2025), achieving statewide canopy goals will require policies that actively target private landowners in disadvantaged communities. Public programs alone can only protect a small portion of the urban forest (Mincey et al., 2012).

### 4.2 Best Model Performance

The statewide fine-tuned model (SFT) consistently outperformed the regionally fine-tuned models, despite the latter’s climate-zone-specific training. This suggests that, for statewide urban canopy mapping, the breadth of annotated training data may matter more than geographic specialization when urban tree structure shares common visual characteristics across regions. Regionally fine-tuned models still performed competitively with fewer annotated tiles, suggesting they may be useful when annotation resources are limited. The LiDAR-trained model showed more spatial offsets and boundary artifacts, likely reflecting misalignment between LiDAR point clouds and NAIP imagery during label generation. Fine-tuning with manually annotated NAIP data reduced these artifacts and generalized well across acquisition years.

### 4.3 The importance of accounting for error in canopy change estimates

Canopy rasters, even when derived from high-resolution imagery and high-accuracy models, are predictions, and without propagating classification error into area estimates, comparisons across years or products can reflect mapping bias rather than true canopy change. This distinction is consequential for California, where AB 2251 mandates a 10% statewide canopy increase by 2035: if baseline and target estimates are both drawn from unadjusted pixel counts, reported progress may not reflect reality. Most large-scale canopy products do not apply error adjustment, instead reporting mapped values alongside classification accuracy metrics as separate quantities. The recent CAL FIRE (2025) statewide urban canopy dataset is a prominent example: mapped canopy cover was reported as 18.85% in 2018 and 18.78% in 2022. When we applied our reference sample to the CAL FIRE product using the Olofsson et al. (2014) framework, error-adjusted estimates increased to 28.15% (95% CI: 5,206-6,106 km²) in 2018 and 23.98% (95% CI: 4,433-5,206 km²) in 2022 –– a difference of nearly ten percentage points in both years (Appendix D). A similar pattern holds for our own dataset, where error-adjusted estimates diverge substantially from mapped pixel totals. The magnitude of this divergence demonstrates that pixel-count totals can systematically misrepresent statewide canopy extent, and that changes in mapped area across years may reflect shifts in classification performance as much as real landscape change. For context, the Chesapeake Bay Land Cover dataset (one of the only other multi-year, large-scale urban canopy change products in the U.S.) reports mapped values with change classification accuracies of 75% (2013-2014) and 56% (2021-2022) without error adjustment (Bouffard et al., 2025), suggesting that this issue extends beyond California.

### 4.4 Comparison with Prior Urban Canopy Mapping Approaches

The statewide canopy cover dataset produced here extends and improves on a growing body of work using deep learning and high-resolution imagery for broad scale urban canopy mapping. Using the subset of reference points overlapping both products, our dataset achieved higher overall accuracy than CAL FIRE (2025) in urban areas (92.2% vs. 86.3% in 2018; 94.0% vs. 88.7% in 2022), with stronger canopy user’s and producer’s accuracies in both years, while additionally covering non-urban CDPs and a longer timeseries. Across the broader literature, U-Net architectures applied to aerial imagery have consistently achieved high accuracy for canopy segmentation. Our SFT model achieves F1 scores of 0.87-0.91 across climate zones at 60 cm resolution over a statewide domain of 39,032 km², exceeding the 79.3% overall accuracy reported by Erker et al. (2019) for a statewide random forest approach in Wisconsin, and comparable to the 91.52% overall accuracy of the UrbanWatch database across 22 U.S. cities (Zhang et al., 2022), while covering a larger and more diverse spatial extent.

### 4.5 Limitations and Future Work

Parallax and imperfect orthorectification can confuse canopy change estimates derived from aerial imagery (Kropp, 2024; Lehrbass & Wang, 2012; Richardson & Moskal, 2014). When acquisition geometry differs across flights or years, the apparent position and footprint of objects can shift, which can inflate or reduce mapped canopy area even when the tree itself is unchanged. Differences in sun elevation between image acquisitions can create shadows that obscure trees differently from year to year. NAIP mosaics vary in orthorectification quality and off-nadir viewing angle both spatially across California and temporally across acquisition years because imagery is collected and processed by multiple contractors and then assembled into a single statewide product. In large metropolitan areas (the Bay Area and Los Angeles Basin), flight lines tend to be denser, which reduces off-nadir effects and generally improves consistency. In many smaller or non-urban regions, flight lines are historically more widely spaced, and off-nadir angles can be larger, increasing geometric distortions (Lillesand et al., 2015).

These effects are not confined only to image seamlines which have higher view angles. Local topography, particularly hillsides, can amplify relief displacement and shape distortions even within a single flight strip, and these terrain-driven effects can vary across years depending on flight geometry and processing (Appendix D). The magnitude of parallax-related bias also depends on canopy form: compact, rounded crowns are less sensitive, whereas tall, conical or vertically layered crowns can have larger footprint changes with larger viewing angles. Together, these factors make it difficult to attribute small year-to-year differences in mapped canopy area solely to true canopy change without explicitly considering acquisition geometry and its interaction with canopy structure. The distortion of canopy area and placement resulting from these effects means that canopy products produced from NAIP imagery should not be used for pixel-to-pixel change comparisons. For that reason, we did not analyze small scale (on each parcel, or by tree) change in urban forest canopy, as they could be the result of changing view angles in any given year. To illustrate the magnitude of this effect, we conducted a small sensitivity analysis by manually digitizing the same 11 tree crowns in NAIP imagery for each acquisition year (2016, 2018, 2020, 2022) in locations where view-angle artifacts were hand selected to be visually apparent (Figure 8). Across years, both the apparent crown footprint and the apparent crown position shifted, producing between year percent area differences that are larger than botanically possible from true tree growth over two-year intervals. While this experiment is not intended to quantify statewide bias, it demonstrates that acquisition geometry and orthorectification can generate canopy change signals at the small scale that are not related to tree growth.

**Figure 8:**
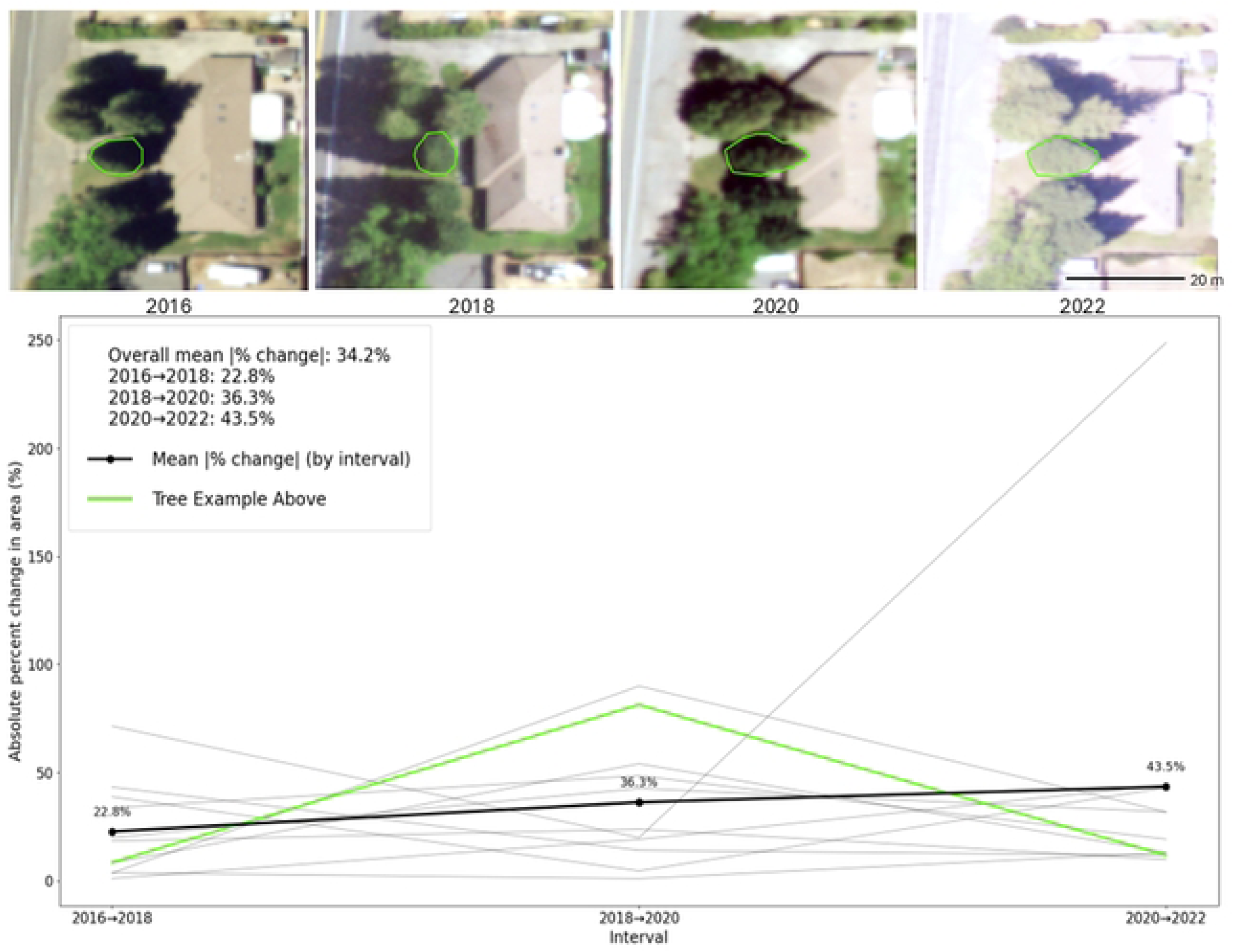
Absolute percent change in canopy area between successive years for 11 manually digitized conical tree crowns with clear view-angle effects. Each gray line shows one tree; the green line corresponds to the tree shown in images above. Changes reflect year-to-year variability in mapped canopy area. Interval means are shown only to indicate the magnitude of variability and should not be interpreted as typical change.

Future work should estimate how frequently these distortions occur and evaluate their contribution to uncertainty in statewide canopy cover and change estimates. In the meantime, these results underscore that NAIP-derived canopy time series contain an inherent error that is unaccounted for in estimates. Policy or management applications that rely on detecting change should emphasize large magnitude changes that are assessed over long time periods, rather than small year-to-year differences that may fall within the range of imaging-related variability. Extending this time series beyond 2022 will further improve the ability to distinguish persistent canopy trends from short-term variability and acquisition artifacts.

## 5. Conclusions

We developed and evaluated a reproducible workflow for generating a multi-temporal, high-resolution canopy cover dataset across California census-designated places using NAIP imagery. The approach combines semi-automated LiDAR-derived training data, manual annotation, and error-adjusted area estimation to support consistent comparisons across years. Statewide canopy cover declined modestly between 2016 and 2022, but the magnitude of change was small relative to total canopy extent and confidence intervals overlapped. These findings suggest that short-term differences in NAIP-derived canopy cover should be interpreted cautiously, especially where image acquisition geometry and orthorectification can introduce apparent change at fine spatial scales. Continued monitoring over longer time periods will improve the ability to distinguish persistent structural trends from short-term variability.

Across most climate zones, canopy cover was relatively stable from 2016 to 2022, with the clearest decline occurring between 2018 and 2020 in the Inland Empire, Inland Valleys, and Southern California Coast. Canopy cover was generally higher in non-urban than urban portions of CDPs, except in the Southwest Desert, Southern California Coast, and Inland Empire, where urban and non-urban patterns were more similar. Most canopy within incorporated urban areas occurred on privately managed parcels, highlighting the importance of engaging private landowners in future canopy planning and protection efforts. Overall, this workflow provides a scalable framework for tracking canopy trends in future NAIP years and can be adapted for urban forest monitoring across other U.S. states.

## 6. Code and data access

Canopy cover data for all four years is open source. Canopy maps can be viewed at: https://experience.arcgis.com/experience/401c3c7a7b3742dcb548178251ed1c91. Code for running and training the model as well as accuracy assessment points are accessible on GitHub at: https://github.com/camipawlak/canopy_cover. Raster data, training chips, and model checkpoints can be downloaded at DOI: 10.5061/dryad.rjdfn2zs9.

## 7. Declaration of generative AI and AI-assisted technologies in the manuscript preparation process

During the preparation of this work AI tools were used to assist with code development, debugging, and minor language editing. After and while using AI, authors reviewed and edited the content as needed and take full responsibility for the content of the published article.

## Appendices

## Appendix A. Inputs and outputs for accuracy assessment and error-adjusted canopy area estimation.

Tables included are:

(A1) Statewide overall by year.

(A2) Urban and non-urban statewide by year.

(A3) Climate zones by year.

Columns report:

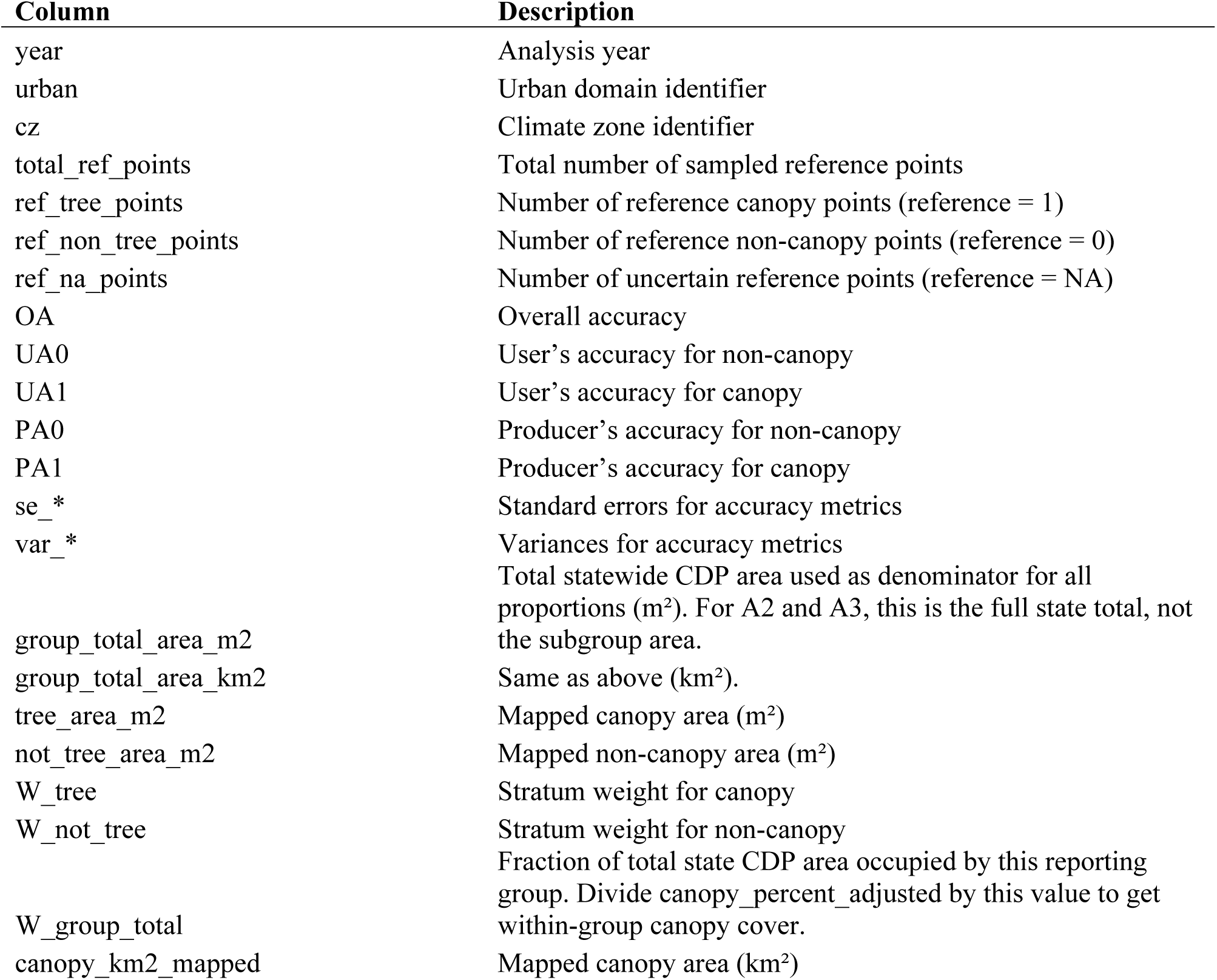
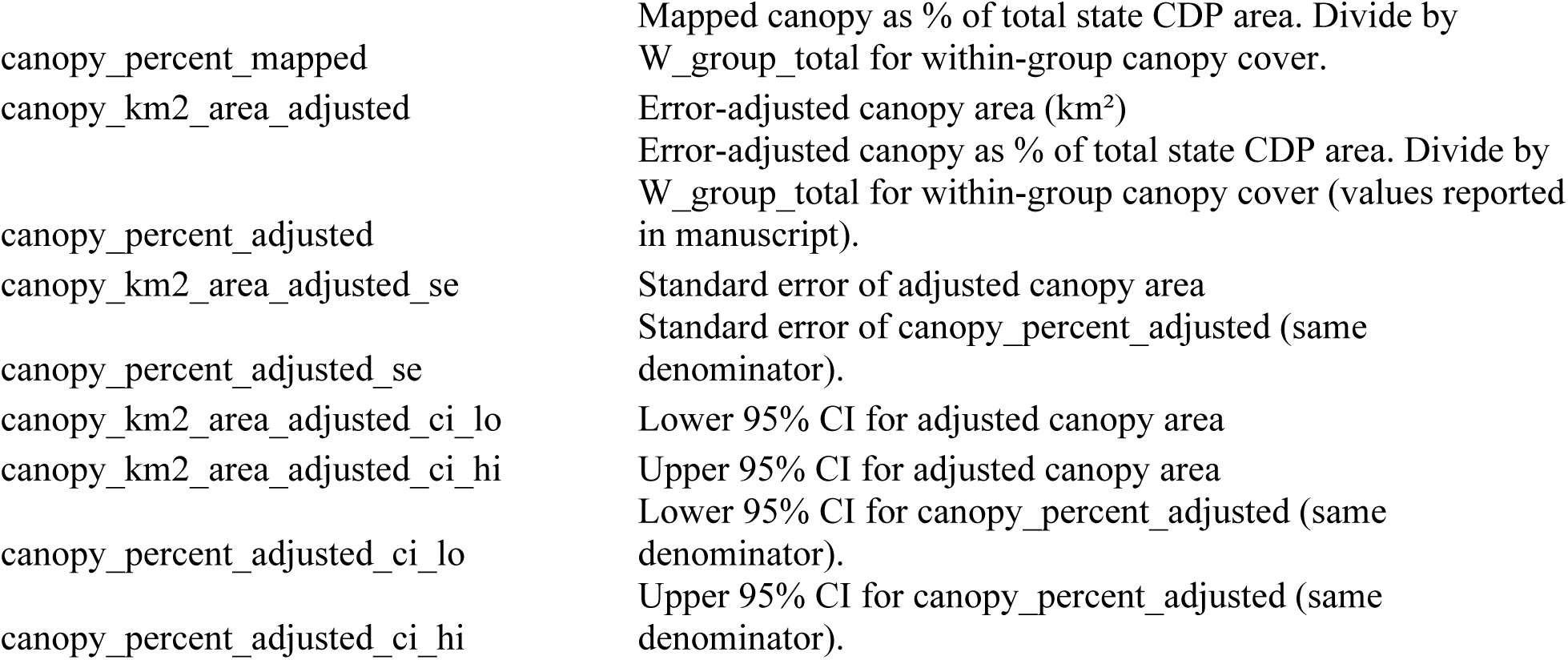

## Appendix B: Inputs and outputs for accuracy assessment and error-adjusted canopy area estimation by year and parcel type (UF_Code). Columns are shared with Appendix A.

## Appendix C. Data sources and cities for LiDAR used to create automated training data.

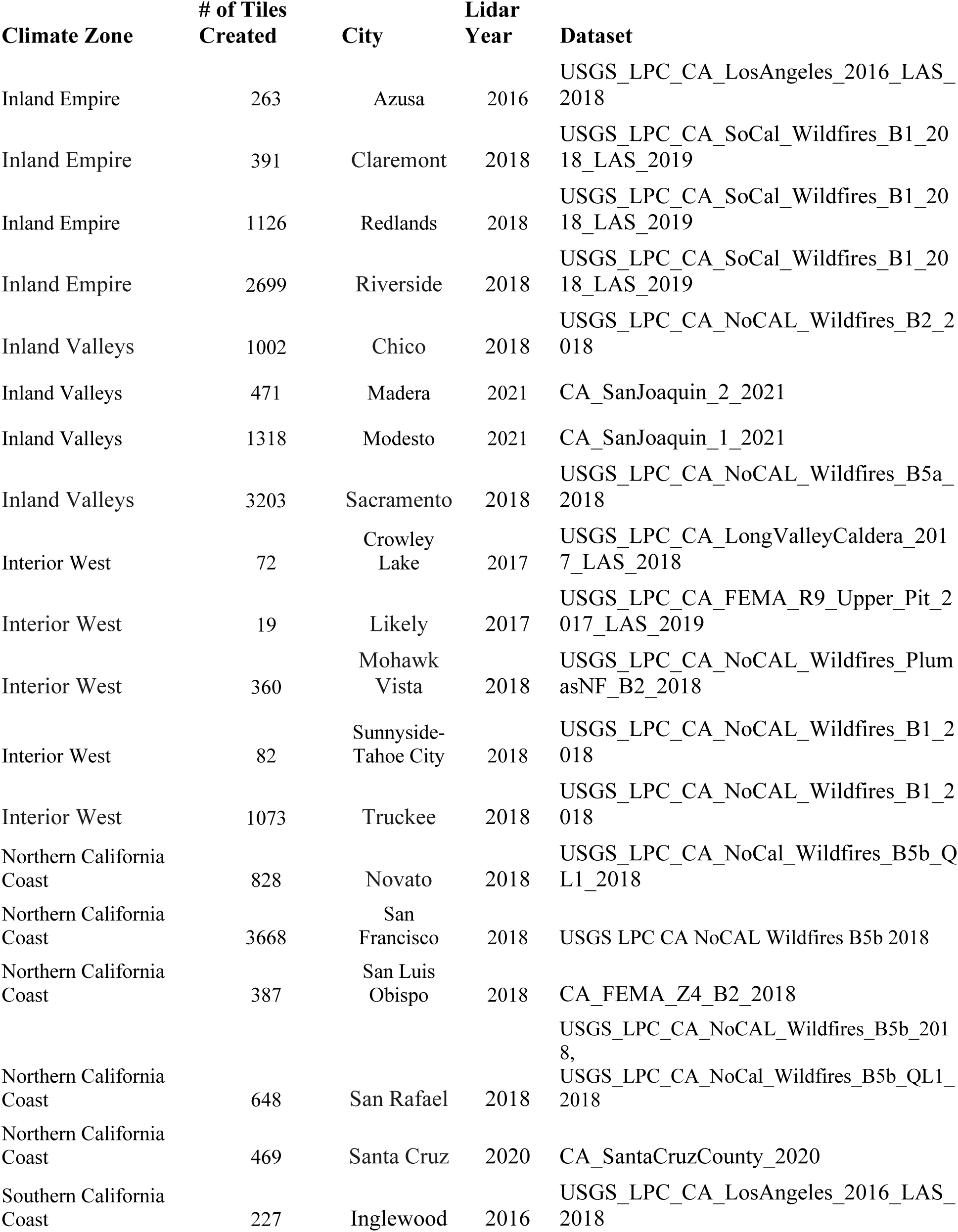
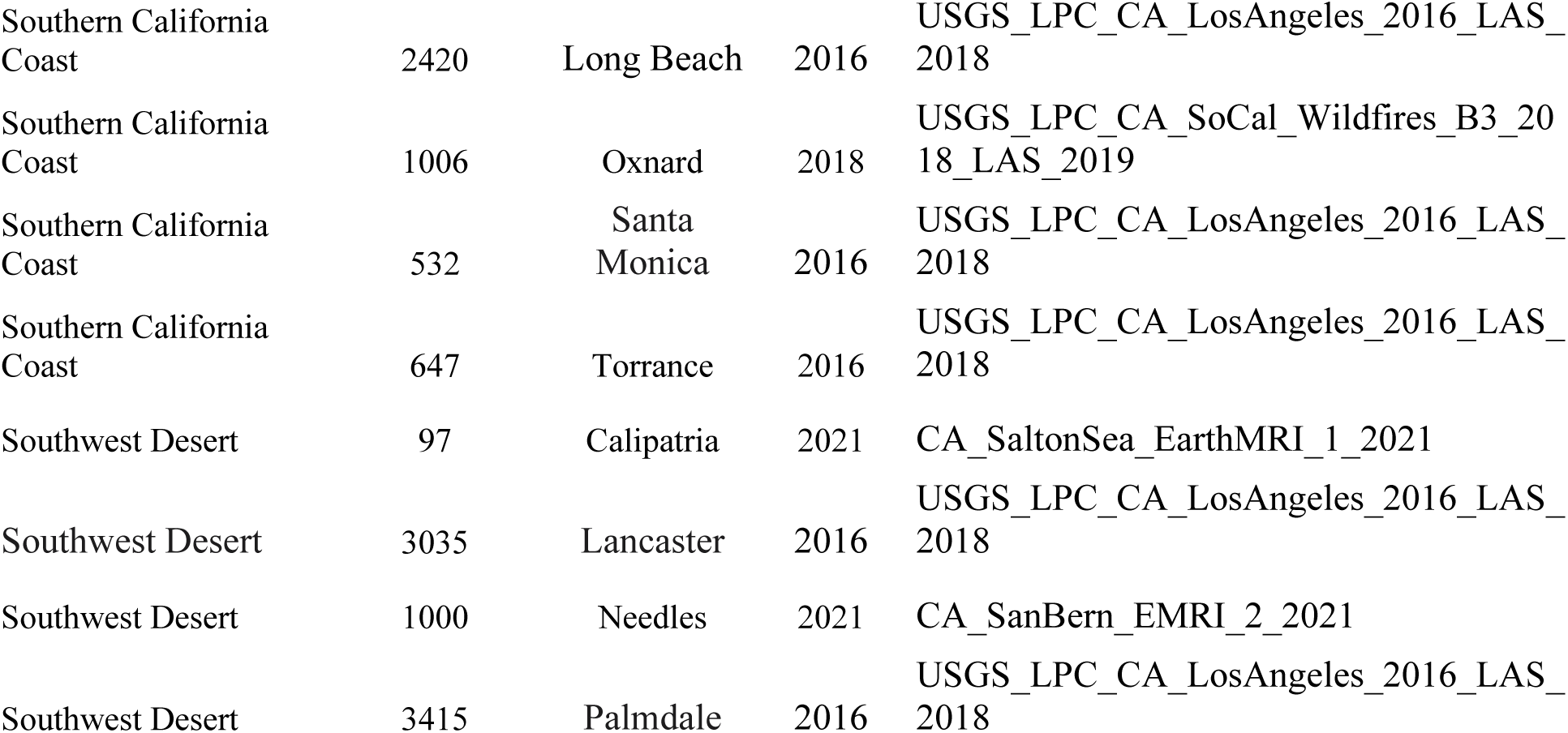

## Appendix D. Examples of relief displacement in NAIP imagery. A) NAIP imagery of trees on a slope, 2018. B) The same trees as A, but in 2020. C) The urban forest with a close to nadir view angle in 2018. D) The same trees as C, off-nadir in 2022.

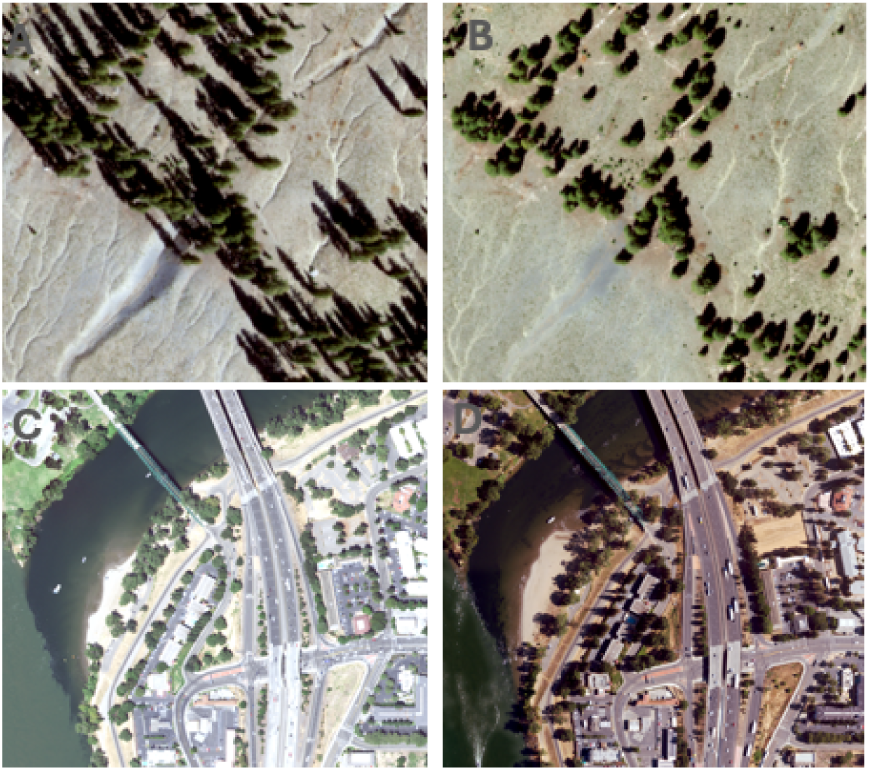

## Acknowledgements

The authors would like to acknowledge the efforts of students who helped create training data for this work: Ryley Chase, Alexandra Crocker, Jessica Baiza, Katharine McDaniel and April Engelmeier. We would also like to acknowledge students who contributed to the LiDAR processing code: Nam Nguyen and Benjamin Hinchliff.

